# Molecular Nearest Neighbors Determine Generalization of Odorant Mixtures

**DOI:** 10.1101/427401

**Authors:** Ryan J. Brackney, Kael Dai, Taleen Der-Ghazarian, Brian H. Smith, Richard C. Gerkin

## Abstract

Most natural odors arise from mixtures of multiple odorants. Some such mixtures are perceived “elementally”, with each odorant component clearly identifiable, while others are perceived “configurally”, with the mixture adopting a perceptual quality distinct from any of the components. While the perceptual similarity of two mixtures is presumably related in some way to the similarity of the corresponding components, given the elemental/configural dichotomy it is unclear if any formal principle can be used to predict mixture similarity. To investigate this problem, we trained mice to respond to a binary reference mixture of structurally similar odorants (*S*^+^) and then tested generalization of this response to other structurally related binary test mixtures. Across 5 experiments, we parametrically varied these mixtures in distinct ways to test candidate models for the perceptual similarity of mixtures. The best-performing model predicted behavioral responses by considering, for each component of the *S*^+^, only the similarity of the most structurally similar (”nearest neighbor”) component of each test mixture. We conclude that for mixture generalization tasks the olfactory system may deemphasize or discard information about mixture components not perceptually “near” enough to any of those in the *S*^+^, consistent with a sparse and elemental rule for perception of structurally-related binary mixtures.

## Introduction

Behavioral evaluation of odor perception in animals is essential to interpreting the neural and molecular bases of olfactory processing. Animals respond to odors innately and through learned behavior shaped by experience of odors with reward. In either case, one experimental approach for evaluating perceptual similarity is to consider a target odorant to which the animal responds, and then to vary some physical quality of the target odorant and determine how strongly the animal generalizes a response to this new test odorant. This approach can establish how perceptually similar the test and target odorants are to each other [Kay et al., 2006, Shepard, 1987, Shettleworth, 2010].

In tests with series of structurally related odorants, the response typically decreases as the test odorant becomes less similar to the target odor; this approach has worked well with monomolecular odorants in both insects and mammals [Abraham et al., 2004, Bhagavan and Smith, 1997, Braun and Marcus, 1969, Cleland et al., 2009, 2002, Duncan et al., 1992, Laska et al., 1999, Linster et al., 2009]. When conditioned to a target odorant, such as an alcohol of a particular carbon-chain length (CCL), animals typically respond most strongly to the conditioned odorant and less strongly as features such as CCL increase or decrease away from the conditioned odorant [Cleland et al., 2002]. Changes in the functional group (e.g. from alcohol to ketone) or its position on the carbon chain can have even stronger effects on generalization responses [Daly and Smith, 2000, Laska et al., 2008, Smith and Menzel, 1989]. In some animals, differential conditioning of two odorants that are close in CCL, where one odorant is reinforced and the other not reinforced, shifts peak responses away from the reinforced odorant and in a direction away from the unreinforced odorant [Daly and Smith, 2000, Fernandez et al., 2009]. This ‘peak shift’ is thought to result from overlapping excitatory and inhibitory gradients distributed along a perceptual (coding) dimension represented in an animal’s sensory system [Mackintosh, 1983].

Consequently, CCL may represent a salient olfactory perceptual dimension, even if not a principal one, or at least part of an ordered series of points in a perceptual state space. That proposition has been also well supported by studies of sensory and early olfactory coding in both insects and mammals [Cleland et al., 2002, Daly et al., 2004, Fernandez et al., 2009]. In the invertebrate olfactory system, binary mixtures evoke neural responses with state space representations that neatly track the ratios of the mixture components [Fernandez et al., 2009]. However, we need more thorough analyses of the ‘metrics’ of olfactory stimulus space to approach a fuller understanding of coding dimensions in the peripheral and central nervous system.

Whether innate or learned, natural odor mixtures typically contain multiple components [Aycı et al., 2005, Aznar et al., 2001, Grosch, 1998, Raguso, 2008]; even semiochemicals, which elicit strong innate responses, such as the urine-based odors used for individual recognition of rodents, can be mixtures of many components [Yamazaki and Beauchamp, 2005]. Some mixtures appear to be perceived as a single ‘configural’ objects [Thomas-Danguin et al., 2014], in which the elements of the mixture cannot be distinguished in the mixture. Other mixtures might be perceived as a sum of individual components in an ‘elemental’ representation [Wagner, 2008]. These odor mixtures can differ based on the composition and/or the ratios among the individual components, which gives rise to many dimensions along which these odors may need to be discriminated.

Because of the potential complexity of natural odor mixtures, it will be necessary to build on the knowledge of responses to mono-molecular odorants by using a step-wise approach to more complex mixtures, beginning with systematically varying simpler, binary mixtures.

Systematically controlled mixtures will allow for investigation of how animals may use different behavioral strategies in evaluating mixtures. The strategies could depend on the nature of the mixture and on the type of reinforcement used to condition a behavioral response. Foraging moths, for example, use only a subset of mixture components from the floral odors they approach for food [Riffell et al., 2014].

Use of the mouse for these types of behavioral studies allows for subsequent genetic manipulation, electrophysiological recording, optical imaging and optogenetic stimulation to test the causal relationships between behavior and events measured in the nervous systems. Mice can typically easily discriminate monomolecular odorants that differ by only a single carbon in CCL [Laska et al., 2008]. Such ‘steep’ generalization makes it difficult to evaluate graded similarities in neural representations. Here we develop a new behavioral protocol for the mouse in which behavioral generalization among mixtures can be evaluated through experimental manipulation. We start with variation in the CCL of a conditioned monomolecular odorant, which serves as a reference for strong discrimination. We then condition to a binary mixture and generate related binary mixtures that differ from the conditioned stimulus in a variety of systematic ways. These mixtures are used to test the role of specific hypotheses about mixture generalization. Finally, we show how the data can be explained by a model in which animals may focus on the use of the most salient ‘predictive’ component of a mixture.

## Methods

### Subjects

Three cohorts (two to four litters in each cohort) of male C57BL/6 (C57) mice (N=42 total, PND 30-60) served as subjects. The mice were single-housed in a colony room with a reverse 12:12 hr light cycle (dark 7 AM to 7 PM). They were water restricted, beginning seven days prior to the experiment. During water restriction, each mouse received 1 mL of water in their home cage at 4:30 pm each day, in addition to any liquid they earned in the experimental session. Experimental sessions occurred between 9 AM and 4 PM, up to seven days a week. All animal use and experimental procedures conformed to guidelines established by the National Institutes of Health (NIH) Guide for the Care and Use of Laboratory Animals and the Arizona State University Institutional Animal Care and Use Committee.

### Apparatus

Experiments were conducted in a “Slotnick-style” 4-channel pinch-valve olfactometer (described in [Slotnick and Restrepo, 2005]) and operant chamber purchased from KNOSYS. All odorants were diluted to 0.1% in mineral oil (Sigma-Aldrich #M3516-1L), except as noted below.

In a previous set of experiments (not shown), we investigated the ability of mice to discriminate between olfactometer channels based on two potential confounded: the sound of the channels, or previously used odorant that might have adsorbed to channel components. We did this by training to criterion (described below) using one odor-containing, rewarded channel, and one blank (air), unrewarded channel. We then tested the same two channels with blank stimuli and a random reward schedule. We found no significant preference for the channel that had previous contained odor, indicating that neither of the two confounds above were a concern in the current experiments.

Experimental events were controlled and recorded by PyOperant (http://bitbucket.org/rgerkin/pyoperant). The reinforcer was approximately 0.006 mL of chocolate milk (Shamrock Farms; Phoenix, AZ) diluted in water (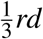 milk by volume) combined with the sounding of a brief (0.3 s) 9 kHz buzzing tone at 65 dB from a set of stereo computer speakers. Subjects were given approximately 0.5 mL of the milk mixture 24 hours prior to the first experimental session to familiarize them with the reward they would experience in behavioral experiments.

### Procedure

The odorant discrimination training and generalization testing protocol was a modified version of the Go/No Go task used by Slotnick and colleagues [Slotnick and Restrepo, 2005]. Each experiment consisted of four phases: Pre-training, Odorant discrimination (full reinforcement), Odorant discrimination (partial reinforcement), and Generalization Testing. Each subject had only one training session per day, consisting of 10 blocks, with each block consisting of 10 trials. Testing sessions occurred on separate days from training sessions.

### Pretraining and Odor Discrimination (Full reinforcement)

Pretraining and Odor discrimination training proceeded as described previously [Slotnick and Restrepo, 2005]. During pre-training, subjects were trained to lick the liquid delivery tube and keep their head in the odor tube for several seconds. No odor was presented during pre-training. Discrimination training consisted of the standard Go/No Go task in which subjects were trained to lick in the presence of one odor, the *S*^+^, and not in the presence of another odor, the *S*−. Each odorant discrimination session began with a “warm up” block of 10 trials in which all trials were *S*^+^ in order to establish robust responding to the *S*^+^ [Nevin and Grace, 2000]. The remainder of the session consisted of 9 blocks of 10 *S*^+^ trials and 10 *S*^−^ trials, ordered pseudo-randomly so that no more than 3 trials of the same type (*S*^+^ or *S*−) occurred consecutively within a block. Once a subject responded with at least 80% correct for two consecutive sessions, it was advanced to partial reinforcement protocol.

### Odorant discrimination (partial reinforcement)

Generalization trials with the novel stimuli were conducted in the absence of reinforcement in order to prevent the subjects from learning new stimulus-reward associations to those novel stimuli. To ensure that subjects would nonetheless persist in responding during the generalization phase, they were trained under low rates of reinforcement during a preceding partial reinforcement phase. Training was identical to odorant discrimination, except the probability of reinforcement on *S*^+^ trials after the “warmup” block was gradually reduced over four sessions. During the first session, 80% of *S*^+^ trials resulted in reinforcement for correct responding. The following three sessions reduced the frequency of reinforcement on *S*^+^ trials to 60%, 40% and then 20%. The next session after 20% reinforcement was Generalization Testing.

### Generalization testing

During generalization testing, the probability that a subject would respond to each of three generalization odorants (*a*, *b*, and *c*) relative to the *S*^+^ was tested. Generalization testing consisted of “warm up” block, as per odorant discrimination, and four generalization blocks. Generalization blocks consisted of eight trials – two trials with each generalization odorant (*a*, *b*, and *c*), and two trials with the *S*^+^. Responding to the generalization odorants was never reinforced. However, to prevent response extinction, responding on one of the two *S*^+^ trials in each block was reinforced, while the other was not. In generalization blocks, trial order was determined pseudorandomly, in that all four types of trials (*S*^+^, *a*, *b*, and *c*) must have occurred before a second trial of the same type occurred.

### Experiments

Five different experiments were conducted across three cohorts of mice. Cohort 1 was trained and tested for experiment 1-5, cohort 2 for experiments 1-3, and cohort 3 for experiment 1. Figure 1 outlines the stimuli used as the *S*^+^ and for generalization testing (*a*, *b*, *c*) in each experiment. For experiments 1 and 5, the stimuli were pure monomolecular odorants. For experiments 2, 3, and 4 all stimuli were binary mixtures that differed in overlap between the *S*^+^ and the test mixtures. Together, these experiments aimed to test different mixture configurations so that any well-fitting model would have to generalize to a large portion of mixture configuration space.

**Figure 1.**
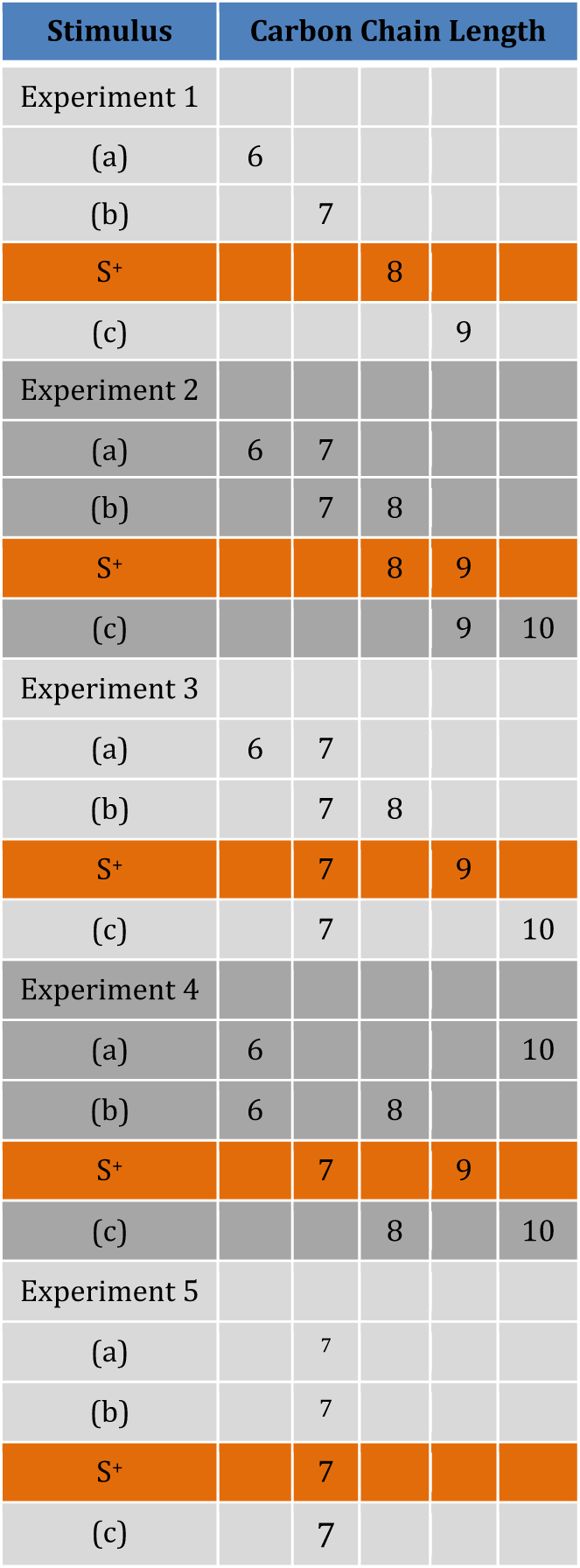
Experimental design. In each experiment type (1-5), each row corresponds to one stimulus, with the *S*^+^ and the three test-only stimuli (a-c) listed for each experiment. For each stimulus, the molecular components are given along the row. Each number corresponds to the carbon-chain length (CCL) *n* of one component, with all components being secondary straight-chain alcohols. For example, a stimulus with only a ”7” refers to 2-heptanol, while one with ”7” and ”8” refers to a binary mixture of heptanol and 2-octanol. All odorants were diluted to 0.1% in mineral oil, except in the vapor pressure control (experiment 5); in that experiment the stimulus is always 2-heptanol, with the dilution (schematized by font size) changing across stimuli. In some experiments with multiple cohorts, the identity of the *S*^+^ and one of the generalization stimuli were switched between cohorts as a robustness check.

1. *1) Single-Linear*. This experiment tested generalization in the limiting case of stimuli with only one component. Each stimulus was a simple dilution of a 2-n-alcohol (where ‘n’ refers to CCL, e.g. *n* = 7 refers to 2-heptanol) to 0.1% concentration by volume in mineral oil. The generalization stimuli (*a*, *b*, *c*) differed from the *S*^+^ by either 1 or 2 carbons in CCL, in either direction.
2. *2) Double-Linear*. This experiment tested the extension of generalization to binary mixtures by shifting the CCL of the test odors while keeping the relative difference between the components the same. Each stimulus was a mixture of a 2-n-alcohol with the corresponding 2-(n+1)-alcohol, e.g. a mixture of 2-hexanol and 2-heptanol. Each component was diluted to 0.05% concentration by volume in mineral oil, so the total volume of alcohol was 0.1% as in the first experiment.
3. *3) Double 1-Shared*. This experiment tested the differential contributions to generalization of shared vs distinct components. Each stimulus was a binary mixture, composed as in Double-Linear. However, each of the generalization stimuli shared one component (2-heptanol) with the *S*^+^, and differed from it in the other component.
4. *4) Double 1-Distant*. This experiment tested the contribution of average CCL to generalization. In other words, can binary mixtures be represented by the mean location of their components in a feature space, with generalization around that mean? Each stimulus was a mixture of 2 odorants, diluted as in the second experiment. However, the generalization stimuli shared no overlapping components with the *S*^+^, but had identical or similar average CCL. For example, the *S*^+^ was a mixture of 2-heptanol and 2-nonanol, with an average CCL of 8, while test stimulus *a* was a mixture of 2-hexanol and 2-decanol, also with an average CCL of 8.
5. *5) Vapor Pressure Control*. Vapor pressure decreases approximately 3.3 fold with each increment of CCL in this molecular series. To ensure that the results in experiments *1*-*4* could not be explained by simple differences in vapor pressures across the molecular series, experiment *5* trained subjects on 2-heptanol at 1.0% dilution as the *S*^+^, and tested them on 2-heptanol at 0.01%, 0.1% and 5.0%.

### Model Types

We modeled the data by considering only the CCL of the molecules composing each stimulus. For a test stimulus *x*, the predicted response probability under all models used the same master equation:

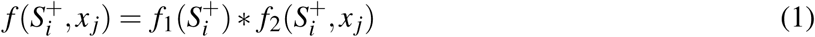

where 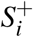 and *x*_*j*_ are the CCL of the *i*^*th*^ and *j*^*th*^ elements of the *S*^+^ and the test stimulus, respectively. The functions are defined as:

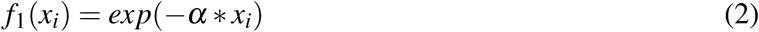

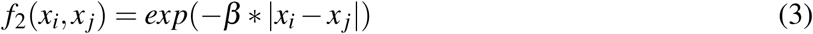

with *α* and *β* free parameters, obtained by maximum likelihood estimation (see below). *f*_1_ represents the intrinsic salience of an *S*^+^ component, and the functional form of *f*_1_ is inspired by the vapor pressure dependence of CCL, which is one component of perceived intensity. *f*_2_ represents the generalization gradient across CCL.

We then considered 3 classes of models (schematized in Fig. 2), varying in computational complexity, i.e. the number of computations of *f* across stimulus components.

**Figure 2.**
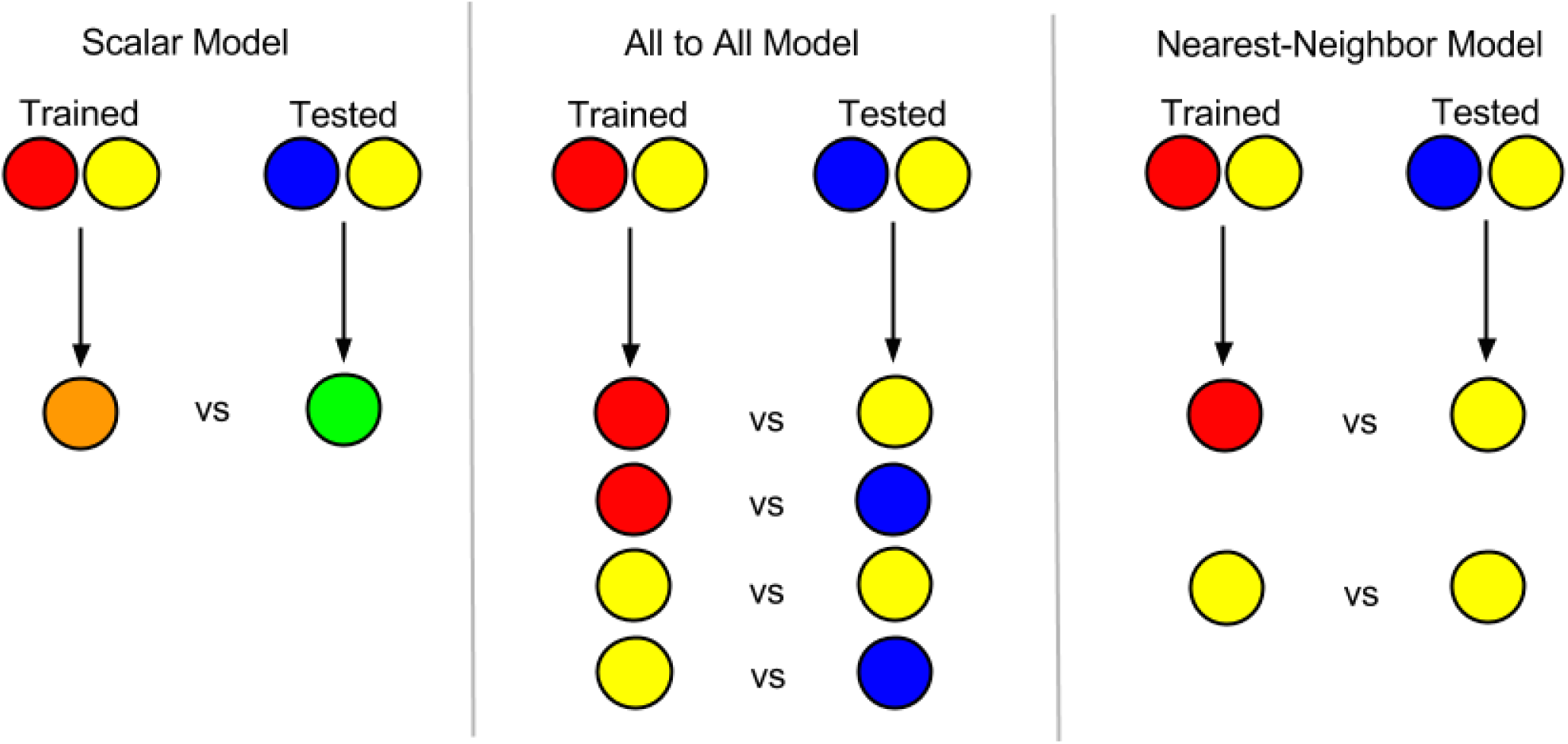
Schematic of three candidate models for olfactory generalization. Circles represents components in a binary mixture, with different colors representing different molecules. Arrows point to possible perceptual representations of stimuli for computation of mixture simularity. (**A**) In the *Scalar* model, each mixture is represented as a single value, average CCL. This value for the trained mixture is compared against the corresponding value for the test mixture, with the similarity of those values determining the probability of response. (**B**) In the *All-to-All* model, each mixture is decomposed into components, and comparison between values is “all to all” between combinations of trained and tested components. All such comparisons are then averaged to determine response probability. (**C**) In the *Nearest-Neighbor* model, each mixture is decomposed, but comparison only occurs between each *S*^+^ component and its “nearest” component (e.g. most similar in CCL) in the test mixture.

1. *Scalar*: Each stimulus is considered as having only one element, represented by the mean CCL of its components. The probability of response (*p*_*r*_) is a function of the difference between the mean CCL of the *S*^+^ and that of the test stimulus, i.e.:

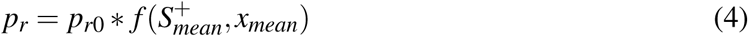
2. *All-to-All*: Each stimulus is represented by *N* = 2 elements, corresponding to the CCL of each component. *p*_*r*_ is a function of the differences between each combination of elements across the *S*^+^ and the test stimulus, summed and normalized.

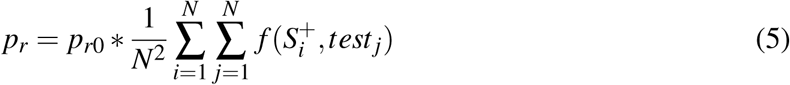
3. *Nearest Neighbor*: Each stimulus is represented by *N* = 2 elements as in *All-to-All*, but *p*_*r*_ is a function only of the difference between each element *i* of the *S*^+^ and the single nearest (component most similar in CCL) element of the test stimulus array, with index *i*†.

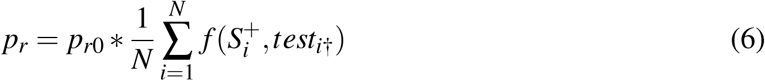

Each model was fit to the pooled data, expressed as response probabilities for each test stimulus in each experiment, using maximum likelihood estimation to obtain the parameters *a*, *b*, and *p*_*r*0_. The first two parameters were each assigned one value shared by all experiments. We allowed *p*_*r*0_ to vary according to the experiment type (although it was still fixed for all subjects and stimuli within each experiment), to account for variable baseline response motivation across experiments. Goodness-of-fit is reported as the mean-squared error between the model fit and the observed response probabilities, and is fundamentally bounded at the low end due to binomial variability (Figure 5, “Noise”).

### Data pooling

The first three experiments were replicated in two to three cohorts of mice (Methods). The identity of the *S*^+^ systematically differed across replications, in order to balance the test stimuli in either direction of CCL. We observed similar results across cohorts and *S*^+^ identities (Figure 3), so we pooled the data for modeling purposes such that one component of the *S*^+^ was designated as the reference component. All other components in the *S*^+^ or the test stimuli were then labeled according to the difference in CCL among their components relative to the reference (Figure 5, dashed lines). For example, for an *S*^+^ consisting of 2-heptanol and 2-octanol, we designated 2-heptanol as the reference, 0, and 2-octanol as +1. Together the *S*^+^ could be labeled (0, +1). A test stimulus of 2-octanol and 2-nonanol is then labeled (+1, +2). This allowed us to present data from experiments using different *S*^+^ identities together on the same plot (Figure 5).

**Figure 3.**
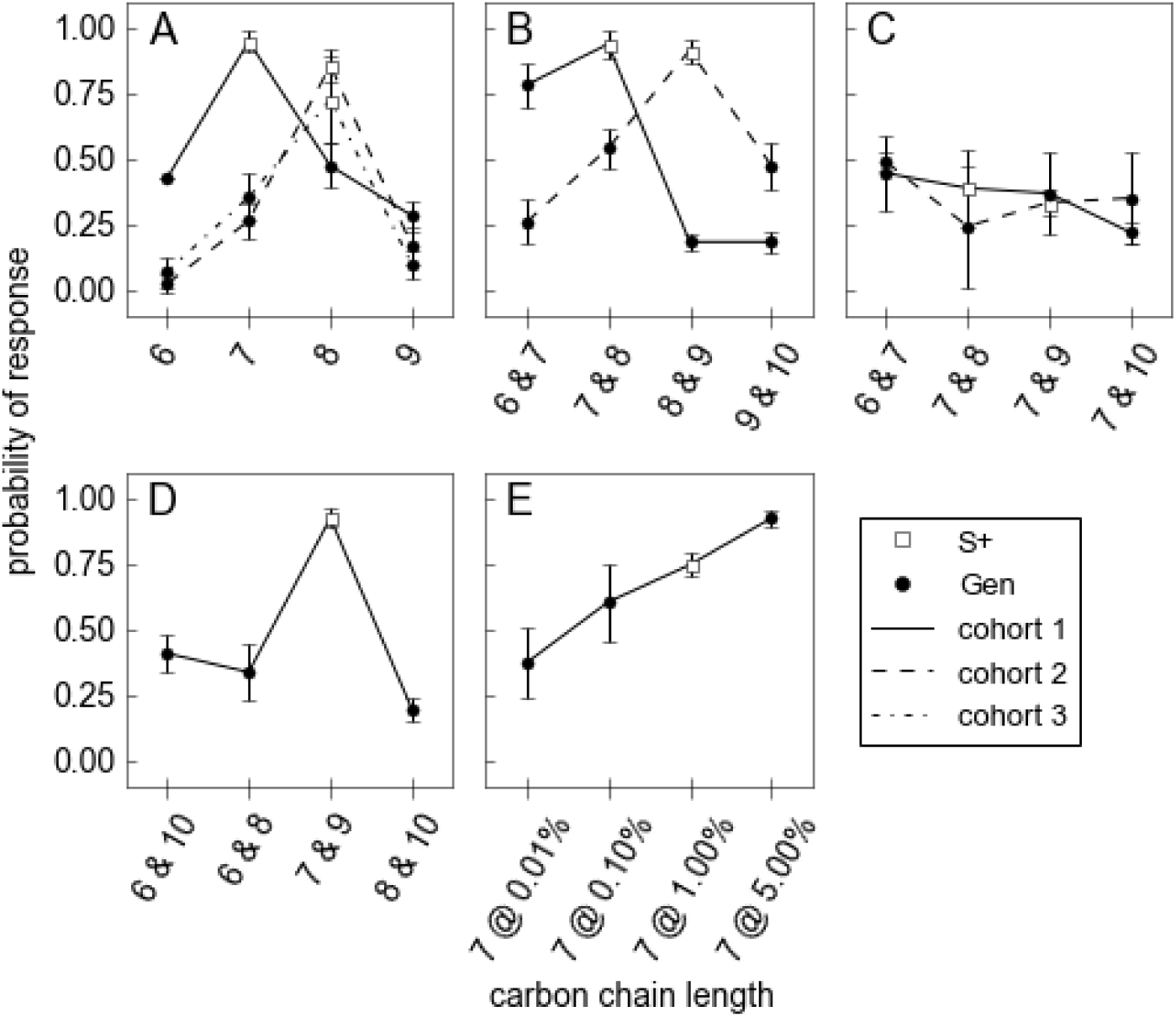
Results of behavioral experiments. Animals were trained to discriminate the *S*^+^ (indicated by an open square) from a background (mineral oil) control. Black circles represent the novel generalization test stimuli (*Gen*, e.g. the a,b and c stimuli for each experiment in Fig. 1) not experienced during training. The Y axis indicates the response probability to each testing and training odor. Error bars indicate the standard error of the mean. Separate lines indicate experiments on three different cohorts of mice. **A**) Exp. 1 Single-Linear: X-axis shows CCL of stimuli, where each stimulus was a secondary straight-chain alcohol, e.g. 2-heptanol. **B**) Exp. 2 Double-Linear: Same as *A*, except stimuli are binary mixtures of molecules at equal concentration. As in *A*, adjacent stimuli are 1-carbon apart in CCL. **C**) Exp. 3 Double-1-Shared: Similar to *B*, except each binary mixture contains 2-heptanol in common. **D**) Exp. 4 Double-1-Distant: Similar to *B*, except each test mixture has the same or similar mean CCL as the *S*^+^. **E**) Exp. 5 Vapor Pressure Control: Test stimuli vary from the training stimulus by concentration, but are otherwise identical. Response probability is a linear function of concentration. n=4 or 5 mice from each cohort used in each experiment.

### Code

All code and analysis is available at http://github.com/quolf/nearest-neighbor.

## Results

### Experiments

In the experiment 1 (“Single-Linear”; Figure 1) we tested the direct molecular similarity between the *S*^+^ and each of the generalization stimuli (*a* − *c*). The results (Fig. 3a) show that for each of three mouse strains generalization responses decreased with increasing CCL differences between the *S*^+^ and the test stimuli, as reported previously for both mammals and insects [Chaudhury et al., 2009, Daly et al., 2001, Laska et al., 2008, Yoder et al., 2014]. Increasing the CCL of the *S*^+^ by one carbon had no significant effect on the shape of the generalization function (Fig. 3a).

In experiment 2 (“Double-Linear”; Figure 1), the *S*^+^ and the test stimuli were a monotonic series of binary mixtures. Generalization to the test stimuli occurred as a function of the degree of overlap of the mixtures with the *S*^+^ (Fig. 3b). However, the test stimuli that contained one component in common with the *S*^+^ (‘b’ and ‘c’ in Fig. 1) elicited higher responses than the corresponding *S*^+^-adjacent test stimuli in experiment 1. This indicates that the generalization gradient is shallower for such binary mixtures than for corresponding monomolecular mixtures (Fig. 3b; gaussian width = 0.75 +/0.02 for experiment 1, 1.09 +/-0.05 for experiment 2). Some generalization asymmetry was observed in one mouse cohort, but this was not significant after a false discovery rate correction for multiple comparisons.

In experiment 3 (“Double-1-Shared”; Figure 1), the test stimuli were binary mixtures that differed by one component from the *S*^+^. In this experiment, generalization was almost totally flat (Fig. 3c), indicating that the identity of the differing component was almost irrelevant. The animals appeared to generalize to the common component, although overall response to the *S*^+^ was lower than observed in the other experiments. We initially suspected that this experiment may have just been a ’failure’, but given that it replicated with a second cohort, that other experiments interleaved with it showed robust responses, and that inspection of the olfactometer revealed no defects, we concluded that this may have just reflected natural variation. A reviewer suggested discarding it, but we concluded that this would be a dangerous precedent in the absence of any overwhelming, fundamental theoretical objection to the experimental outcome.

In experiment 4 (“Double-1-Distant”; Figure 1), the two components of each generalization test stimulus were distinct from those of the *S*^+^, but they spanned a similar range of CCL or had the same mean CCL. Only a low level of generalization was observed (Fig. 3D), with a narrow generalization width (0.66 +/-0.05) similar to that observed in the ‘Single-Linear’ experiment.

In experiment 5 (“Vapor Pressure Control”; Figure 1) the test stimuli were identical to the *S*^+^ (2-heptanol), except they all varied in concentration. Alcohols of lower molecular weight tend to have higher vapor pressures. Therefore, in experiments 1-4 generalization could have been driven primarily by similarity to the partial pressure of the vapor phase of the *S*^+^. In some cases, different vapor phase concentrations of an odorant can produce qualitatively different percepts, so it is possible that a subject’s response probability might not be monotonic in concentration. To control for this possibility, we performed an experiment using a single odorant (2-heptanol), with the test stimuli differing from the *S*^+^ only in concentration. We found that response probability was approximately linear across concentration, with no deviation reflecting a specific preference for the concentration of the *S*^+^. Had vapor pressure of the molecule been responsible for a controlling perceptual feature of the odor, the response probability would have peaked at the concentration corresponding to the *S*^+^. Therefore differences in vapor pressure did not principally drive response probability differences in experiments 1-4.

### Modeling

To quantitatively account for the generalization observed in each of the five experiments, we considered three potential approaches for computing the perceptual similarity of mixtures, and then fit a simple model for each. These models shared a similar core feature–simple functions of absolute and relative CCL–but differed in which sets of mixture components are involved (Figure 2). Each model contained 3 parameters: the baseline response probability (*p*_*r*_0), the salience of individual components as expected from e.g. differences in the partial pressure of the vapor phase (due to differences in vapor pressure) (*α*), and the steepness of generalization for components differing in CCL (*β*). Figure 2 qualitatively illustrates the set of computations in the three different models, and a quantitative description is provided in the Methods. The code used to implement the computations can be found at http://github.com/quolf/nearest-neighbor.

Computational complexity varied across the three models (Figure 2). It was simplest in the “*Scalar*” model, consisting only of a single comparison of average CCL between the *S*^+^ and the test mixture. It was most complex in the “*All-to-All*” model, consisting of *N*^2^ comparisons, one for each pair of components across the *S*^+^ and the test mixture (each having *N* = 2 components). The “*Nearest-Neighbor*” model was intermediate in complexity; it consisted of only *N* comparisons, one for each component in the *S*^+^ and its corresponding ‘nearest’ component in the test mixture.

### Model fits

We estimated the parameters for each model that maximized the likelihood of the observed response probabilities, using the same parameters (for each model) across experiments and cohorts. Figure 4 shows the average mean squared error across all five experiments. While the the *Scalar* model showed the poorest fit, the *All-to-All* and *Nearest-Neighbor* models fit the data better, with the latter yielding the lowest mean squared error (Fig. 4). Identical rank-ordering of results was obtained by evaluating the likelihood function directly (not shown). While the difference between the *All-to-All* and *Nearest-Neighbor* models was not statistically significant, maximum likelihood estimation of model parameters indicated that the observed data was ∼ 6*x* as likely under the *Nearest-Neighbor* model than under the *All-to-All* model, and ∼ 110*x* more likely than under the *Scalar* model.

**Figure 4.**
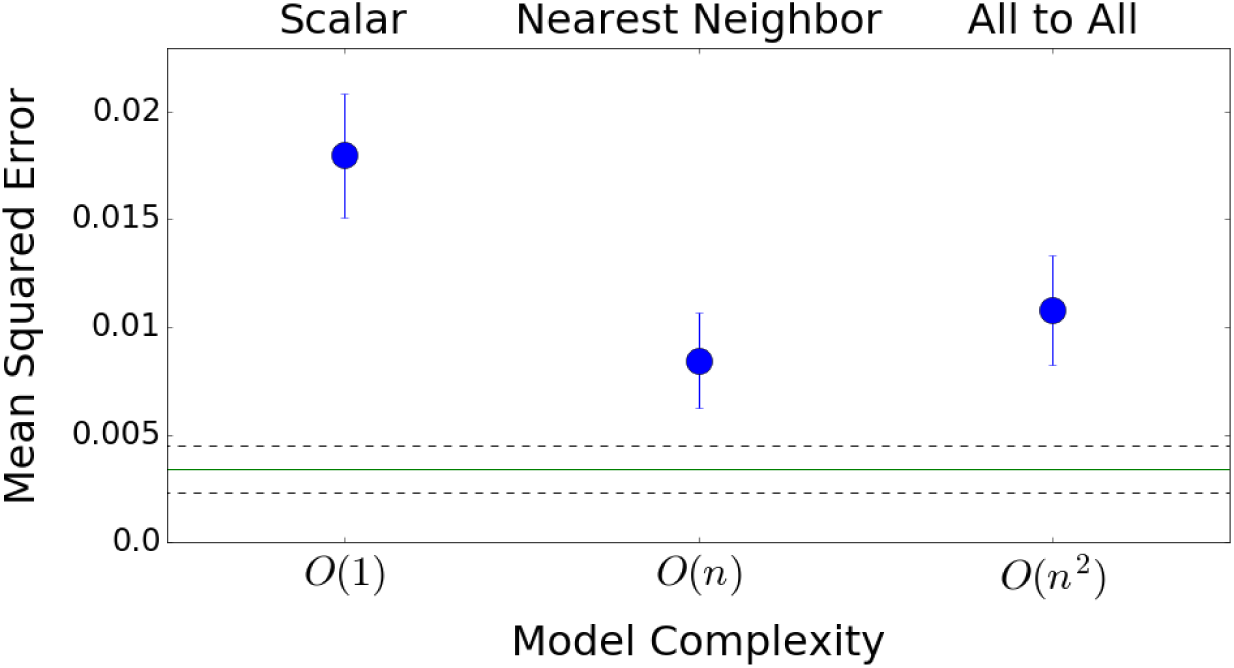
Mean-squared error (MSE) for each model fit. Models were fit to all experiments using the same parameters, except for baseline response probability which was allowed to vary across the five experiments. Error bars reflect the standard deviation of MSE, derived from 100 synthetic datasets using the observed response probabilities and binomially distributed numbers of responses per experiment. The green line represents the average MSE expected from simple binomial sampling variability, i.e. from a finite number of Bernoulli trials, and reflects the theoretical limit of model accuracy. The dashed lines reflect +/-1 standard deviation of this MSE, which was calculated by drawing new Bernoulli trials from the observed mean response probabilities for each animal for each stimulus for each experiment. The *Nearest-Neighbor* model yielded the best fit of the three models, although it was not significantly more accurate than the more computationally complex *All-to-All* model.

**Figure 5.**
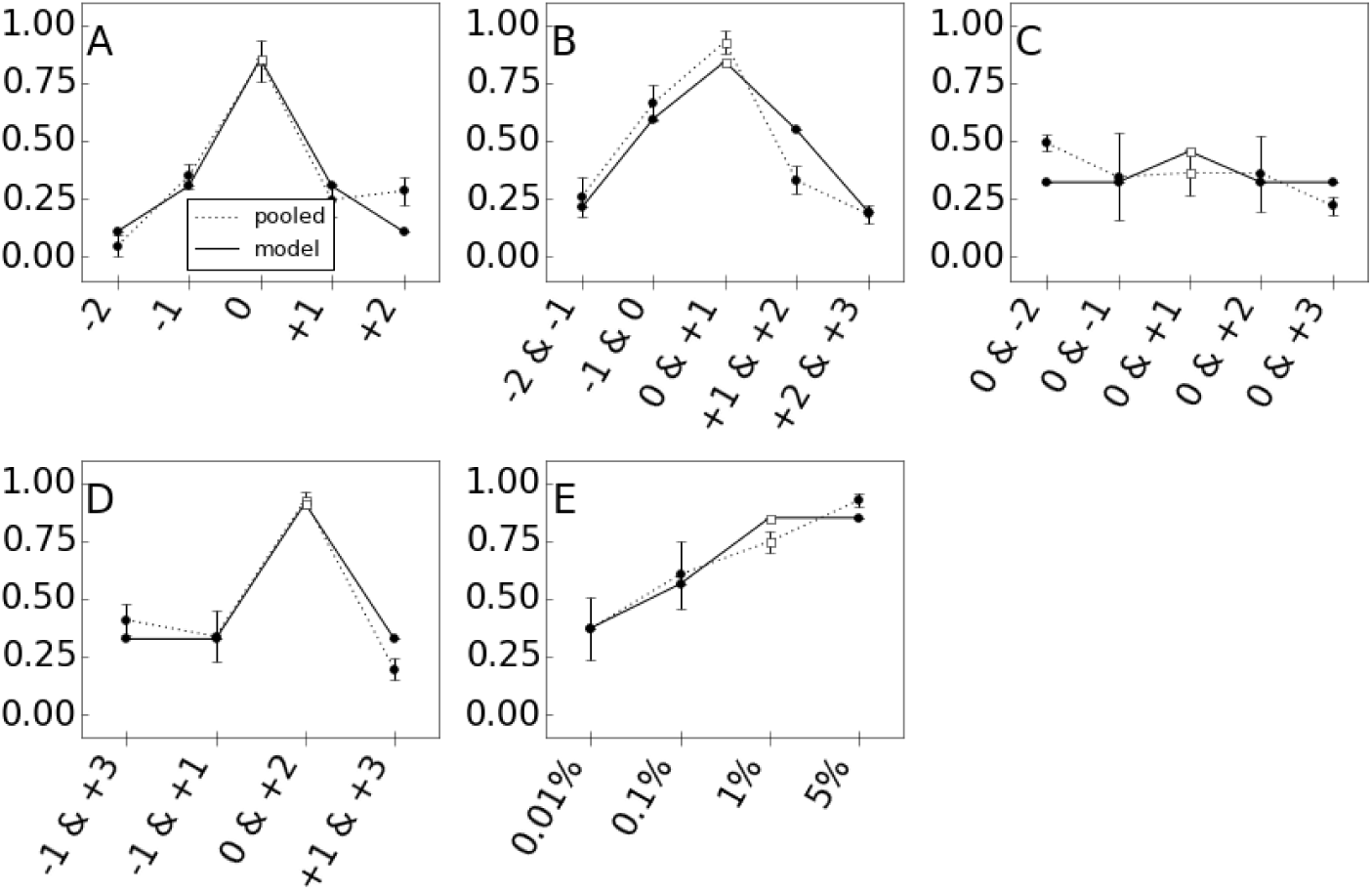
Model fits for the *Nearest-Neighbor* model. For each experiment in Fig. 3, the data were normalized so that the first component of the *S*^+^ is defined to have a relative CCL of 0. Each stimulus is then described according to the CCL of its components relative to the *S*^+^. The resulting “pooled” experimental results are then averaged across mouse strains (dotted line). The solid line gives the prediction from the *Nearest-Neighbor* model for each experiment. The same parameters were used across fits to the three models (Fig. 2), except for the baseline response probability, which was allowed to vary by experiment. This baseline response probability allowed us to account for the variability in responsiveness across experiments, whether it resulted from random or systematic variation. To the extent that experiment 3 showed weak responses, the baseline response probability parameter effectively gives this experiment a smaller weight in the estimation of other model parameters.

Even with a perfect model, some deviation between model and data is expected due to binomial variability, i.e. the finite number of trials. We assessed this by generating synthetic data using the same observed response rates to each stimulus, with the same number of trials. The deviation between the observed and the synthetic data (averaged across 1000 synthetic data sets) reflects the lower bound of model fit error, and thus a theoretical upper bound of model performance. The *Nearest-Neighbor* approaches this theoretical upper bound, suggesting that it is unlikely that a much better model can be constructed, at least with so few parameters. When each experiment is considered separately, the *Nearest-Neighbor* had the lowest mean squared error (of the three models) for four out of the five experiments. Fig. 5 shows model fits for the *Nearest-Neighbor* model to the pooled data.

## Discussion

In order to understand how perceptual similarity is determined across mixtures, we conducted five olfactory generalization experiments with binary mixtures of straight-chain aliphatic alcohols. In each experiment, we trained mice to associate a single mixture (*S*^+^) with reward and then tested the generalization of their conditioned responses during exposure to the *S*^+^ or to related mixtures. Across experiments, the relationship between the *S*^+^ test stimuli was varied to determine what features of the *S*^+^ were most effectively generalized during testing.

While generalization was poor for single molecule stimuli (Figure 5a), as expected, binary mixtures of the same stimuli were sufficient to produce generalization of the *S*^+^ (Figure 5b-d). We developed a family of models based on possible interactions between molecular components of the *S*^+^ and ensuing test stimuli, and fit them to the data from all five experiments. The *Mean* model reflected one type of ‘configural’ strategy for olfactory identification, assigning a single value to each mixture, reflecting the mean CCL of the components. That model predicts that increasing the CCL of one mixture component can be offset by decreasing the CCL of another. This model was a poor fit to the data, and we were unable to devise a variant with better performance that also relied on representing each binary mixture with a single scalar value (not shown). The *All-to-All* model was a much better fit, suggesting that there is still some ‘elemental’ information in each mixture being utilized by the animals in their perceptual decisions. However, despite using more information than the *Nearest-Neighbor* model, the *All-to-All* model did not perform better, and in fact performed worse (though not significantly so), suggesting that some mixture components may be irrelevant to generalization, and that including them in a model calculation simply adds noise to response predictions. This is surprising because the complexity of the task could have required a model with many more degrees of freedom, but instead was well-fit by the *Nearest-Neighbor* model, which had relatively few. It is admittedly difficult to imagine a prior, biologically-inspired case for the *Nearest-Neighbor* (how would the olfactory system truly extract elements and then implement such a comparison?). However, typically, when exploring the space of statistical models, implementations with small number of parameters that nonetheless nearly saturate fit quality can illustrate important general principles about how a system can be expected to operate. The *Nearest-Neighbor* model was remarkably simple and effective, suggesting that an important predictor of generalization may be the similarity of the *most similar component* of the test mixture to a given component of the *S*^+^.

How might this principle apply with more complex mixtures, containing far more than 2 components? We note that, unlike the other model variants, the *Nearest-Neighbor* model has the interesting property of reducing to a ‘fractional overlap’ rule as the number of components becomes large, as observed in mixture discriminability studies in humans [Bushdid et al., 2014] and other mammals [Romagny et al., 2015]. In other words, for large *N*, the same equations predict that the response probability will be roughly determined by the fraction of components in the test stimulus which are identical to components in the *S*^+^. This is true even though the model operates without giving special preference to identical components, simply because identical components, if present, will naturally be nearest-neighbors. Although generalization is a distinct phenomenon from discrimination [Cleland et al., 2009], this may nonetheless suggest that the ‘fractional overlap’ rule is simply a large *N* generalization of a more basic theoretical principle of elemental coding. Indeed, evidence for elemental coding in complex mixtures is observed in human imaging data [Howard and Gottfried, 2014]. Alternatively, there might be a strategy-switch between elemental and configural approaches to mixture perception that depends on task demands, including mixture complexity [Chandra and Smith, 1998, Coureaud et al., 2011].

CCL is an intuitive molecular feature but this does not mean that the olfactory system makes any use of it. The most predictive models of olfactory perception instead use complex features from computational chemistry software that may better map onto patterns of olfactory receptor activation [Keller et al., 2016, Snitz et al., 2013]. Indeed, elemental rules for mixture generalization may break down with sufficient complex mixtures, as nonlinear effects of and interactions between component concentrations may make simple elemental concentration scaling rules inapplicable. This may explain why qualitative differences in odor character at varying concentrations are more common in complex than in simple mixtures.

One recent study predicted perceptual similarity of mixtures using the ‘angle difference’ of feature vectors in a high-dimensional space of such molecular features [Snitz et al., 2013]. Applying their model (with parameters fit from human data on discrimination) to the mouse data we acquired resulted (unsurprisingly) in a poor fit to our mouse generalization data (not shown). With a sufficiently large corpus of species- and paradigm-specific data, this model might be more successful in predicting generalization using complex mixtures.

We used CCL as the only molecular feature of interest; whether similar results might be observed for variation in functional group, hydrophilicity, molecular weight, or some other feature is a question that ambitious future experiments can address. Because CCL is so easily parametrically controlled, and because we observed generalization even for mixtures with only a 1-carbon difference in CCL, we are optimistic about the use of CCL in future experiments with additional degrees of freedom. For example, components ratios could also be modified, since even binary mixtures with 100-fold ratios between components are perceived differently in rodents than the corresponding pure odorants [Yoder et al., 2015]. Such investigations could be paired with recording or imaging of the olfactory system in order to understand the neurophysiological bases of odor generalization and possible implementations of any of the rules discussed above.

## Acknowledgements

We acknowledge awards from the Arizona Alzheimer’s Consortium and the Army Office of Scientific Research, which funded this research.

